# Co-Infection of cattle in Virginia with *Theileria orientalis ikeda* genotype and *Anaplasma marginale*

**DOI:** 10.1101/2021.04.28.441839

**Authors:** Vanessa J. Oakes, S. Michelle Todd, Amanda A. Carbonello, Pawel Michalak, Kevin K. Lahmers

## Abstract

*Theileria orientalis ikeda* is a newly identified agent of bovine infectious anemia in the United States. Although it is transmitted by separate tick hosts than *Anaplasma marginale* – a bacterial etiology of bovine infectious anemia –the geographic distributions of these two infectious organisms overlap, with co-infection reported in some cattle. Only anaplasmosis has approved effective treatment in the United States. To provide rapid diagnostic information for producers with anemic animals, we developed a duplex qPCR for *A. marginale* and *T. orientalis*. With a cut-off of 38 cycles, the duplex assay has a sensitivity of 96.97% and a specificity of 100% for *A. marginale*; with a cut-off of 45 cycles, the duplex assay has a sensitivity and a specificity of 100% for *T. orientalis*. In addition to providing a tool for improved clinical decision-making for veterinarians and producers, this qPCR facilitates the study of co-infection rate of cattle in Virginia. Of 1,359 blood samples analyzed, 174 were positive for the presence of *T. orientalis*, 125 were positive for the presence of *A. marginale*, and 12 samples were positive for both *T. orientalis* and *A. marginale.* This indicated that co-infection of both of these etiologies of bovine infectious anemia does occur within the state of Virginia. It is likely that this pattern of infection will be seen in regions where *T. orientalis* and *A. marginale* are endemic, despite the difference in tick vectors.

## Introduction

The *Theileria orientalis* complex describes several genotypes of a species of non-proliferative, theilerial hemoprotozoan. Within the complex, the *ikeda* and *chitose* genotypes (*T. orientalis ikeda* and *T. orientalis chitose,* respectively) are capable of causing disease. *T. orientalis ikeda* in particular is increasingly implicated as a causative agent of bovine infectious anemia in the United States, a vector-borne disease characterized by hemolytic anemia, icterus, general malaise, ill thrift, and sporadic abortions. Although rarely fatal, affected animals are often poorly producing; consequently, this is a disease of economic importance in the countries in which it is found.^9^ *T. orientalis* is most effectively transmitted by the Ixodidae tick *Haemaphysalis longicornis,*^8^ a tick which has recently been discovered in several states along the US Eastern seaboard.^13^ *T. orientalis ikeda* has recently been identified in cattle in Virginia, affecting animals that were also parasitized with *H. longicornis* ticks;^10^ this tick has been confirmed as a competent vector for the Virginia *T. orientalis ikeda*.^7^ The animals that were clinically affected in this outbreak presented with the typical signs of anemia, icterus, and general malaise. In Virginia, this clinical presentation is identical to the blood infection caused by the bacterium *Anaplasma marginale. Haemaphysalis longicornis* is not capable of transmitting *A. marginale,*^5^ and in many other parts of the world, the geographic distribution of *T. orientalis ikeda* and *A. marginale* do not overlap. In Virginia, however, the initial outbreak of *T. orientalis ikeda* occurred in areas where anaplasmosis has been diagnosed historically, suggesting an overlap in geographic distribution in this region. Further, the predicted range of *H. longicornis* based on modeling suggests its presence in areas of the country where anaplasmosis is routine.^12^ An animal included in the initial outbreak study in Virginia was positive for both *T. orientalis ikeda* and *A. marginale* by conventional PCR and Sanger sequencing^10^. As a bacterial infection, anaplasmosis is a treatable condition using oxytetracycline, a cost-effective drug approved for use in food animals. Theileriosis is markedly more difficult to treat, and no effective drugs are approved for use in food animals in the United States, although Buparvaquone in Australia has been effective.^4^ Therefore, making an early distinction between anaplasmosis and theileriosis is critical for immediate and effective clinical decision-making that has welfare, productivity, and economic implications. It is important that the tools used to assist in this process are cost-effective, to be of use to producers.

In response, we developed a duplex real-time PCR assay that is sensitive for both *A. marginale* and *T. orientalis*. We further identify the genotype of *T. orientalis* for *Theileria*-positive samples with an additional multiplex qPCR genotype assay (included in the cost to the producer). Previous multiplex assays examining *Theileria* spp., *Babesia* spp. Protozoan, and *Anaplasma* spp. bacteria have been developed in ruminants,^1,6^ so we sought to fill the niche for cattle.

## Materials and Methods

### Blood

Whole blood samples were submitted from client-owned cattle across the state of Virginia. Animals represented clinical submissions by private veterinarians, as well as herds owned and maintained by the Virginia Department of Corrections (VADOC), and as part of an ongoing, random sampling surveillance effort of animals sent to auction by the Virginia Department of Agriculture and Consumer Services (VDACS). Whole blood was collected in purple-top BD Vacutainer® blood collection tubes containing EDTA anticoagulant (Becton, Dickinson and Company). Except for animals submitted by referring veterinarians as part of a routine clinical diagnostic workup, all animals were randomly sampled, and the presence and degree of clinical signs were unknown. In total, 1,359 blood samples were available for evaluation.

### DNA Extraction

DNA was extracted from EDTA anticoagulated blood with the DNeasy® Blood and Tissue Kit (Qiagen) following the manufacturer’s protocol with a few modifications.^10^ The initial blood volume for DNA extraction was 100μL, and Applied Biosystems™ VetMax™ Xeno™ Internal Positive Control DNA (ThermoFisher Scientific) was added to the lysis buffer (Buffer AL) at 20,000 copies per sample. DNA was eluted twice in 50 μL of Buffer AE pre-heated to 56°C for a total elution volume of 100μL.

### Duplex qPCR

The duplex assay is a TaqMan-based assay that utilizes primers and probes designed to detect the major surface protein 1b (msp1b) gene of *A. marginale*, and the major piroplasm surface protein (MPSP) of *T. orientalis*. The primers and probes for MPSP are sensitive for *T. orientalis* but are not genotype-specific; to further characterize genotype, a second assay is run on those samples that return positive for *T. orientalis*.

Amplification for the msp1b and MPSP were accomplished in the same reaction. The duplex qPCR reaction consisted of Applied Biosystems^®^ TaqMan™ Environmental Master Mix 2.0 (ThermoFisher Scientific), 300nM of each *T. orientalis* forward and reverse primers, 600nM of each *A. marginale* forward and reverse primers, 100nM of *T. orientalis* Universal probe, 200nM of *A. marginale* probe, 0.8μL of Xeno™ VIC™ Primer-Probe Mix (ThermoFisher Scientific), and 2μL of DNA template in a 20μL reaction. The primers and probes for *T. orientalis* and *A. marginale* have been previously published.^2, 3^ Fluorophores and quenchers were altered from the published probes in order to allow for appropriate multiplexing. (Table 1) The probes utilize three separate fluorescent tags - FAM™ for *A. marginale*, NED™ for *T. orientalis*, and VIC^®^ for the internal positive control. Amplification was completed in an Applied Biosystems™ 7500 Fast Real-time PCR System (ThermoFisher) using standard cycling and the following run method: (95°C ^10:00^) (95°C ^0:15^, 60°C ^1:00^)_x45_. Reactions were completed in Applied Biosystems^®^ MicroAmp^®^ Fast 8-Tube Strips with Applied Biosystems^®^ MicroAmp^®^ Optical 8-Cap Strips (ThermoFisher Scientific). A sample lacking DNA template was included as a negative control with each run. A *T. orientalis* known positive sample, an *A. marginale* known positive sample, and 2,000 copies of VetMax™ Xeno™ Internal Positive Control DNA (ThermoFisher Scientific) were included in separate reactions with each run to serve as positive controls for the respective fluorophore channels.

**Table 1.**
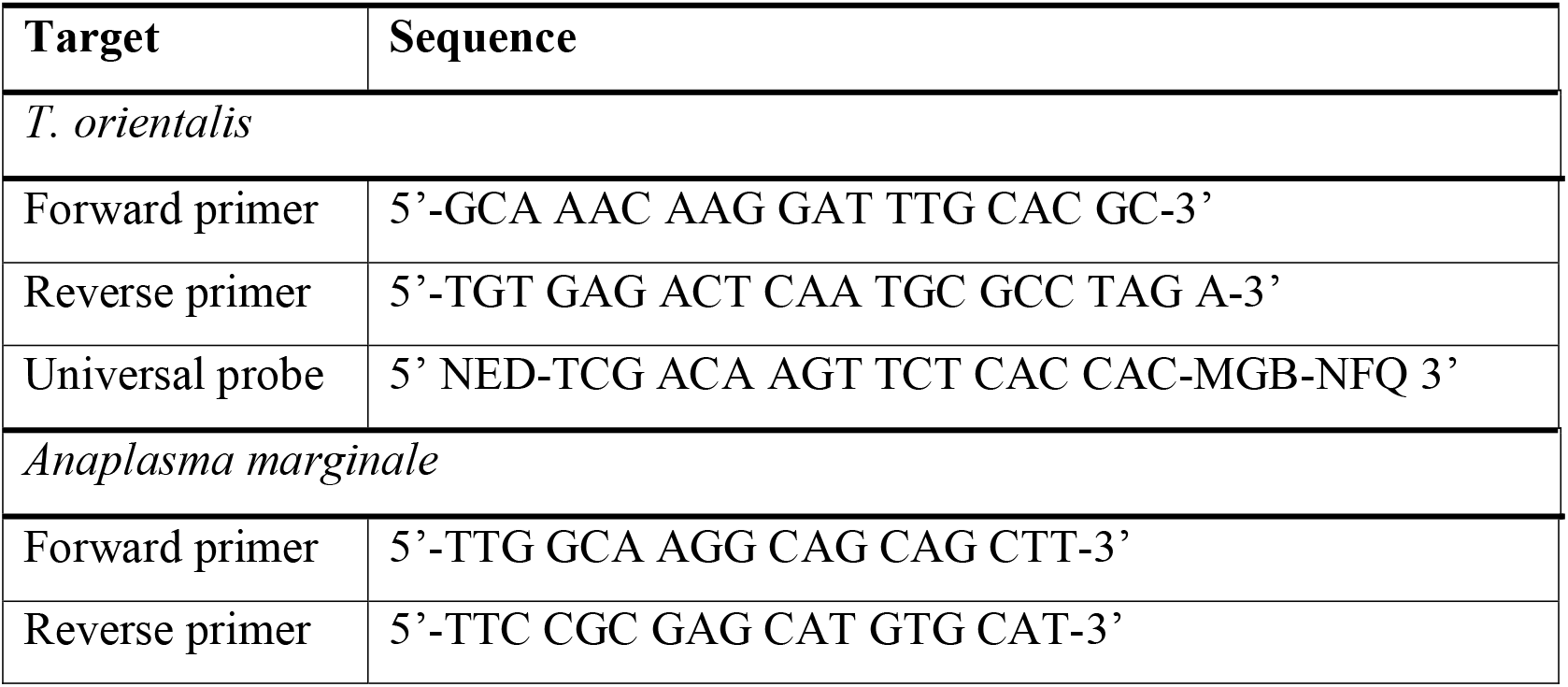
Primer and probe sequences used for *T. orientalis* and *A. marginale* duplex PCR assay.

### Validation: Limit of Detection

To determine the limit of detection for *T. orientalis*, three separate identical qPCR runs were completed per the previously described protocol. Eight, 10-fold serial dilutions of gBlocks™ Gene Fragments (Integrated DNA Technologies) with the universal MPSP gene sequence capable of detecting each of the genotypes *ikeda, chitose* and *buffeli* were used. The limit of detection was determined as the lowest dilution at which samples on all three plates were positive, and a limit of detection was determined for each of the three *T. orientalis* genotypes of interest. Similarly, the limit of detection was determined for *A. marginale* using serial dilutions of an *A. marginale*-positive blood sample confirmed positive by external qPCR and shown to have 1.22% infected RBC.

### Validation: Repeatability

Inter-assay and intra-assay repeatability studies were conducted. Intra-assay repeatability tests the precision of a single run by examining the high, medium, and low concentrations of reference DNA in quintuplicate in one run on one day. In this assay, the average Ct is compared to the standard deviation of the five replicates of a given concentration. Inter-assay repeatability tests high, medium, and low concentrations of reference DNA in quintuplicate on each of six consecutive days. In this assay, the average Ct values of a given concentration are compared across the 6 consecutive plates. All repetitions were performed on the same Applied Biosystems^®^ 7500 Fast Real-Time PCR system (ThermoFisher Scientific).

### Validation: Sensitivity and Specificity

The sensitivity and specificity of the *T. orientalis* and *A. marginale* targets within the duplex assay were determined with receiver operating characteristic (ROC) statistics. Our genotype-specific *T. orientalis* qPCR described below was used as the gold-standard test, against which the duplex qPCR results were compared. For *A. marginale*, qPCR was completed either at the United States Department of Agriculture’s Agricultural Research Service, Animal Disease Research Unit (USDA-ARS-ADRU) (Pullman, WA, USA) or at the Washington Animal Disease Diagnostic Laboratory (WADDL) (Washington State University, Pullman, WA, USA) as the gold-standard test.

### Genotyping qPCR

A second multiplex assay that utilizes distinct probes specific for the MPSP of three of the major *T. orientalis* genotypes - *chitose, ikeda*, and *buffeli* further characterizes the genotype. The forward and reverse primers are the same as those used for *T. orientalis* in the duplex qPCR protocol above, but with the use of a distinct set of probes to distinguish between the genotypes.^2^ (Table 2)

**Table 2.**
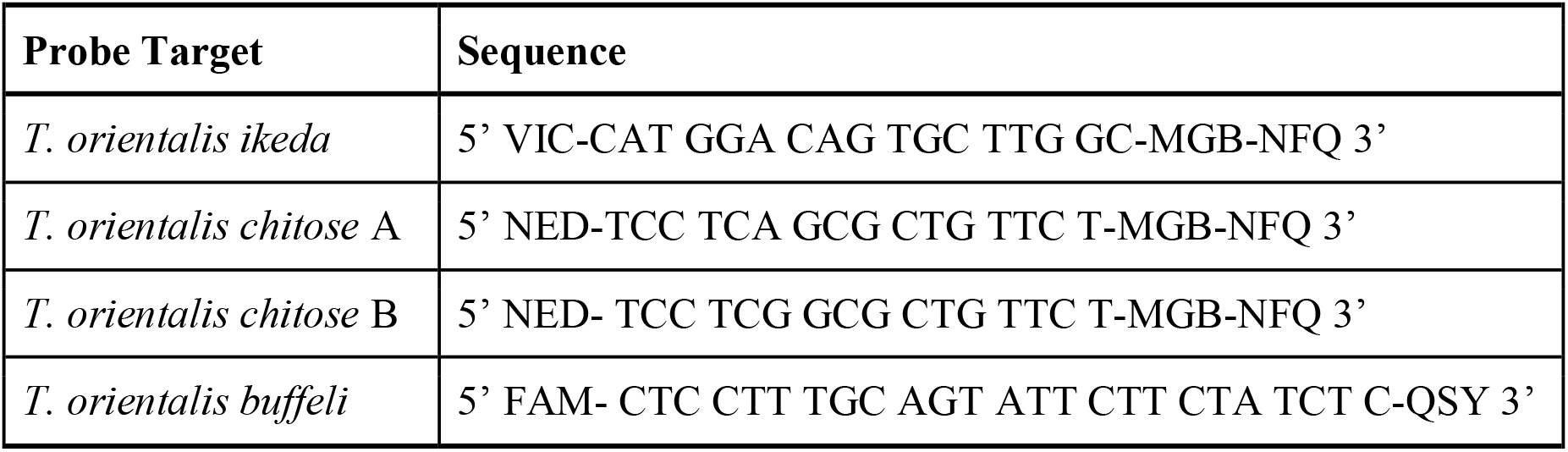
Genotype-specific probe sequences used for the *T. orientalis* genotyping assay.

DNA extracted from whole blood samples that were positive for *T. orientalis* on the duplex qPCR were used for the genotype multiplex qPCR. The 20 μL reactions consisted of Applied Biosystems^®^ TaqMan™ Environmental Master Mix 2.0 (ThermoFisher Scientific), 300nM each *T. orientalis* forward and reverse primers, 250nM of Ikeda probe, 100nM of Chitose A probe, 150nM Chitose B probe, 100nM Buffeli probe, and 2μL of DNA (Table 2).The instrumentation, run method, and plastics were the same as for the duplex qPCR Assay. A sample lacking DNA template was included with each run as a negative control. Positive controls for each of the three genotypes were included in each run.

### Chitose Conventional PCR and Sanger

DNA that tested positive for the *chitose* genotype of *T. orientalis* in the duplex qPCR were subjected to two rounds of conventional PCR (cPCR) targeting the MPSP and SSU genes of *T. orientalis*. The reaction for the first round of cPCR consisted of Invitrogen™ Platinum^®^ PCR SuperMix High Fidelity (Thermo Fisher Scientific), 400nM each of sense and anti-sense primers, and 10.5μl DNA in a 25μl reaction volume. Primers (Integrated DNA Technologies) for MPSP were 5’-CTTTGCCTAGGATACTTCCT-3’ (sense) and 5’-ACGGCAAGTGGTGAGAACT-3’ (anti-sense).^11^ Primers for SSU were 5’-ATTGGAGGGCAAGTCTGGTG-3’ (sense) and 5’-CTCTCGGCCAAGGATAAACTCG-3’ (anti-sense).^9^ The PCR program was as follows: 95°C ^2:00^ (95°C ^0:15^, 57°C ^0:30^, 72°C ^1:00^)_x29_, 72°C 7:00, 12°C ^0:00^. A second round of cPCR was carried out using the amplicons from the first round as template using the same reaction components and PCR program. Second round amplicons were electrophoresed on a 1% TBE-agarose:EtBr gel and visualized under UV light. DNA bands were purified from the gel using the QIAquick® Gel Extraction Kit (Qiagen) and submitted to the Fralin Life Sciences Institute, Genomic Sequencing Center for Sanger sequencing.

## Results

### Duplex qPCR

1,359 individual DNA samples were available for qPCR testing. 186 (13.7%) were positive for the presence of *T. orientalis*, and 137 (10.1%) were positive for the presence of *A. marginale*. Included in these numbers are 12 samples (0.88%) that were positive for both *T. orientalis* and *A. marginale*.

### Limit of Detection

The limit of detection for *T. orientalis* genotypes *ikeda* and *buffeli* were 1×10^8^ picomoles. For the *chitose* genotype, the limit of detection was 1×10^9^ picomoles. *A. marginale* was detected according to dilution rate, given the nature of the known positive sample. The limit of detection of *A. marginale* was a 10^−5^ dilution.

### Repeatability

As calculated, there was significant variability between intra- and inter-assay repetitions of the *A. marginale* component of the assay. However, the variability did not impact the ultimate interpretation of the test results; negative results remained negative, and positive results remained positive. Table 3.

**Table 3.** Comparison of One-Way ANOVA results following the repeatability assay.

| Target | Concentration | One-Way ANOVA | Variance |
| --- | --- | --- | --- |
| *A. marginale* | Undiluted | <0.0001 | Yes |
|  | 10<sup>-2</sup> | 0.0025 | Yes |
|  | 10<sup>-4</sup> | 0.0007 | Yes |
| *T. orientalis buffeli* | Undiluted | 0.7828 | No |
|  | 10<sup>-1</sup> | 0.9730 | No |
|  | 10<sup>-3</sup> | 0.5484 | No |
| *T. orientalis chitose* | Undiluted | 0.2784 | No |
|  | 10<sup>-1</sup> | 0.4041 | No |
|  | 10<sup>-2</sup> | 0.4091 | No |
| *T. orientalis ikeda* | Undiluted | 0.1583 | No |
|  | 10<sup>-1</sup> | 0.7370 | No |
|  | 10<sup>-2</sup> | 0.1495 | No |

### Sensitivity and Specificity

For *A. marginale*, the assay sensitivity was 96.97% and the specificity was 100% when a cut-off of 38 cycles was applied. (Table 4, Table 5) For *T. orientalis,* both the sensitivity and the specificity were 100% when a cut-off of 45 cycles was applied (the final cycle of the qPCR program). (Table 6, Table 7) The gold standard utilized for *A. marginale* was an external qPCR, run either at the United States Department of Agriculture’s Agricultural Research Service, Animal Disease Research Unit (Pullman, WA, USA), or at the Washington Animal Disease Diagnostic Laboratory (Washington State University, Pullman, WA, USA).

**Table 4.**
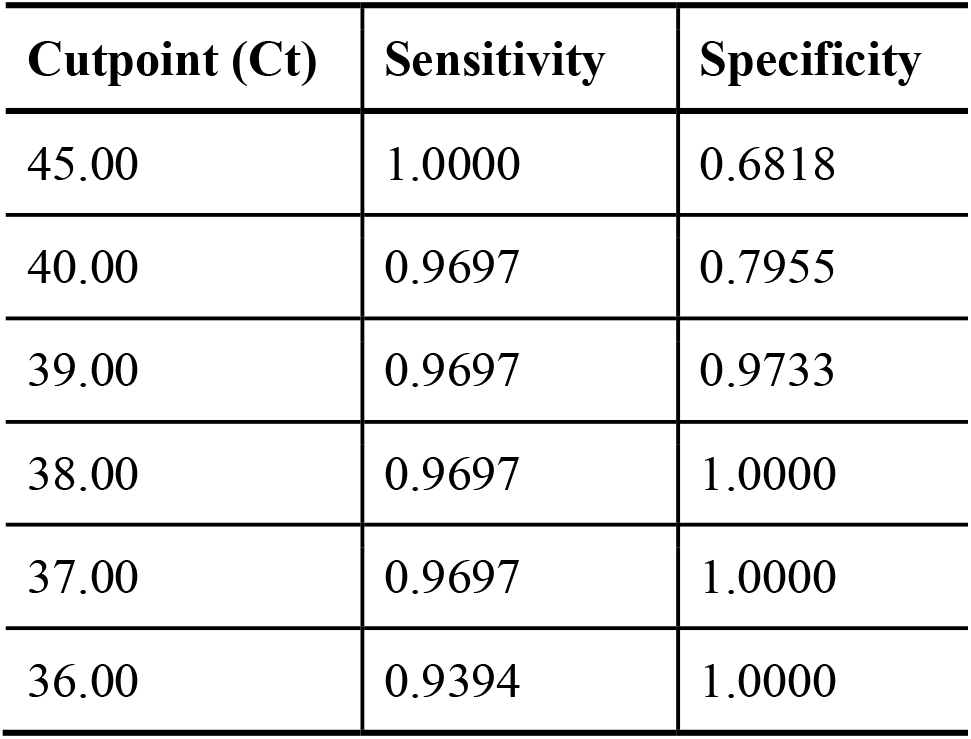
A comparison of the sensitivity and specificity values for *A. marginale* when compared to an external qPCR gold standard.

| Cutpoint (Ct) | Sensitivity | Specificity |
| --- | --- | --- |
| 45.00 | 1.0000 | 0.6818 |
| 40.00 | 0.9697 | 0.7955 |
| 39.00 | 0.9697 | 0.9733 |
| 38.00 | 0.9697 | 1.0000 |
| 37.00 | 0.9697 | 1.0000 |
| 36.00 | 0.9394 | 1.0000 |

**Table 5.** An overview of the positive and negative cases for *A. marginale* when compared to an external qPCR gold standard.

| **Cutpoint 38.00** |  |
| --- | --- |
| True positive | 32 |
| False positive | 0 |
| False negative | 1 |
| True negative | 44 |
| Total | 77 |

**Table 6.** A comparison of the sensitivity and specificity values for *T. orientalis* when compared to a validated qPCR gold standard.

| **Cutpoint (Ct)** | **Sensitivity** | **Specificity** |
| --- | --- | --- |
| 45.00 | 1.0000 | 1.0000 |
| 40.00 | 1.0000 | 1.0000 |
| 39.00 | 1.0000 | 1.0000 |
| 39.00 | 0.9697 | 1.0000 |
| 37.00 | 0.9388 | 1.0000 |
| 36.00 | 0.9 | 1.0000 |

**Table 7.** An overview of the positive and negative cases for *T. orientalis* when compared to a validated qPCR gold standard.

| **Cutpoint 45.00** |  |
| --- | --- |
| True positive | 49 |
| False positive | 0 |
| False negative | 0 |
| True negative | 34 |
| Total | 83 |

The gold standard utilized for *T. orientalis* was the *T. orientalis* genotype qPCR developed by the Virginia Tech Animal Laboratory Services (Virginia-Maryland College of Veterinary Medicine), described below.

### Genotyping

Because of the use of a universal probe for the MPSP gene of *T. orientalis*, an additional genotyping assay was required for those samples that tested positive for *T. orientalis* in the duplex assay. The genotype multiplex assay was run using extracted DNA, applying the same *T. orientalis* forward and reverse primers, but utilizing genotype-specific MPSP probes. Samples of known genotype were included as positive controls. Out of 186 *T. orientalis*-positive samples from the duplex PCR, 159 of them (85.5%) were consistent with the *ikeda* genotype, 21 were consistent with the *chitose* genotype, and the remaining 6 (3.2%) were undetermined. None were consistent with *buffeli*. There were no mixed infections with multiple genotypes.

## Discussion

*T. orientalis ikeda* genotype/genotype 2 is an emerging pathogen within Virginia, and is increasingly implicated globally as an etiologic agent of bovine infectious anemia. The clinical signs of theileriosis and anaplasmosis are identical, consisting of anemia, icterus, poor-doing, decreased productivity, and occasional, sporadic abortions. However, in the United States, oxytetracycline is an approved and effective drug for the treatment of *A. marginale*, whereas there exists no approved drug for the treatment of *T. orientalis ikeda*. Consequently, there is a need for an affordable, effective diagnostic test capable of differentiating between theileriosis and anaplasmosis.

Theileriosis and anaplasmosis overlap within Virginia. This is most likely attributable to a mixed tick infestation in the region. In the United States, *Dermacentor* spp. ticks are the vectors for *A. marginale.*^14^ While *Dermacentor nuttali* ticks have been shown to transmit some genotypes of *T. orientalis* in some parts of the world,^15^ there is no evidence that they are competent vectors for the *ikeda* genotype specifically. Similarly, *H. longicornis*, the primary tick vector of *T. orientalis ikeda* globally, is not a competent vector for *A. marginale.*^15^

Also of interest in this current study is the presence of *T. orientalis chitose* in some of the submitted clinical samples, without co-infection with *ikeda* or *A. marginale*. Like *ikeda*, this is a genotype of emerging importance, as it is increasingly implicated as a non-transforming, pathogenic disease of cattle.^9^ Current understanding on the *T. orientalis* complex suggest *ikeda* and *chitose* have the potential to be pathogenic; *buffeli* is endemic to the United States and elsewhere in the world, and is an incidental, non-pathogenic finding.^9,10^ This difference in genotype pathogenicity is the rationale behind the genotyping component of the assay described in this paper.

The high sensitivity for *A. marginale* and *T. orientalis* makes this duplex qPCR assay a useful screening tool for producers in Virginia and in other localities with co-localization of anaplasmosis and theileriosis. Further investigation into the geographical distribution of *A. marginale* and pathogenic genotypes of *T. orientalis*, including *chitose,* as a probable agent of disease is warranted.

## Acknowledgements

The authors would like to extend immense gratitude to the veterinarians and producers involved in ongoing surveillance efforts.

